# Non-invasive brain-spine interface: continuous brain control of trans-spinal magnetic stimulation using EEG

**DOI:** 10.1101/2020.08.10.230912

**Authors:** Ainhoa Insausti-Delgado, Eduardo López-Larraz, Yukio Nishimura, Ulf Ziemann, Ander Ramos-Murguialday

**Author notes:** These authors contributed equally: Ainhoa Insausti-Delgado and Eduardo López-Larraz. Correspondence: Ainhoa Insausti-Delgado, Institute of Medical Psychology and Behavioral Neurobiology, University of Tübingen, Silcherstr. 5, 72076, Tübingen, Germany, /77501.

## Abstract

Brain-controlled neuromodulation therapies have emerged as a promising tool to promote functional recovery in patients with motor disabilities. This neuromodulatory strategy is exploited by brain-machine interfaces and could be used for restoring lower limb muscle activity or alleviating gait deficits. Towards a non-invasive approach for leg neurorehabilitation, we present a set-up that combines acquisition of electroencephalographic (EEG) activity to volitionally control trans-spinal magnetic stimulation (ts-MS). We engineered, for the first time, a non-invasive brain-spine interface (BSI) to contingently connect motor cortical activation during leg motor imagery with the activation of leg muscles via ts-MS. This novel brain-controlled stimulation was validated with 10 healthy participants who underwent one session including different ts-MS conditions. After a short screening of their cortical activation during lower limb motor imagery, the participants used the closed-loop system at different stimulation intensities and scored system usability and comfort. We demonstrate the efficiency and robustness of the developed system to remove online stimulation artifacts from EEG regardless of ts-MS intensity used. All the participants reported absence of pain due to ts-MS and good usability. Our results also revealed that ts-MS controlled afferent and efferent intensity-dependent modulation of the nervous system. The here presented system represents a novel non-invasive means to neuromodulate peripheral nerve activity of lower limb using brain-controlled spinal stimulation.

## 1 Introduction

Spinal neural networks are in charge of generating locomotor patterns and have the capacity to modulate them even in the absence of brain control (Dimitrijevic et al., 1998; Edgerton and Roy, 2012; Hultborn and Nielsen, 2007). Targeting these self-regulated circuits, also known as central pattern generators (CPGs), for restoring lower limb impairments is the goal of spinal neuromodulation approaches (Minassian et al., 2017; Taccola et al., 2018). During the last decades, invasive electrical stimulation of spinal neuronal pools has been investigated in animals (Alam et al., 2017; Gerasimenko et al., 2008; Ichiyama et al., 2005) and spinal cord injury (SCI) patients (Formento et al., 2018; Gill et al., 2018; Grahn et al., 2017; Harkema et al., 2011) to restore gait patterns.

In this line, magnetic stimulation of the spinal cord presents an alternative modality to neuromodulate the spinal networks non-invasively (Nardone et al., 2015b). In clinical environments, non-invasive magnetic stimulation has been widely used to investigate the central and peripheral nervous systems (Groppa et al., 2012; Nardone et al., 2015a; Rossini et al., 2015). Magnetic stimulation at the spinal level can activate peripheral motor axons at their exit from the spinal cord, evoking muscle action potentials (Knikou, 2013; Matsumoto et al., 2013; Ugawa et al., 1989). Given this knowledge, several studies evidenced the feasibility of repetitive trans-spinal magnetic stimulation (ts-MS) to induce locomotor rhythms using open-loop (Gerasimenko et al., 2010) or EMG-triggered protocols (Nakao et al., 2015; Sasada et al., 2014). However, these procedures lacked volitional and natural brain control.

Neural interfaces allow transferring volitional neural commands between different neuronal populations, bypassing the damaged pathways (Jackson and Zimmermann, 2012). Using brain activity to control the direct stimulation of the spinal cord below the injury level is a natural manner of mimicking the flow of the descending commands from the brain to the spine. This phenomenon has motivated the development of brain-spine interfaces (BSIs) that aim at artificially connecting brain and spinal neural networks to recover motor function (Bonizzato et al., 2018; Borton et al., 2014; Zimmermann and Jackson, 2014). The BSIs record neural activity of the brain reflecting motor intentions and transform this activity into commands for spinal stimulation (Alam et al., 2016; Capogrosso et al., 2018, 2016; Yadav et al., 2020). These neural signatures associated with motor execution or motor attempt can be also detected even in patients with motor deficits (López-Larraz et al., 2018a, 2015), which makes them suitable for BSI control. In order to favor Hebbian plasticity and promote functional recovery, a timely linked brain activity encoding motor intention and peripheral afferent neural activation is essential (Kato et al., 2019; López-Larraz et al., 2018b; Mrachacz-Kersting et al., 2016; Nishimura et al., 2013a; Ramos-Murguialday et al., 2013). In BSIs, peripheral neural activity is generated by the spinal stimulation, modulating the excitability of spinal networks (Hofstoetter et al., 2018; Hubli et al., 2013) and generating muscular contractions of the limbs (Gerasimenko et al., 2018).

To date, BSIs have only been developed as implantable systems, and tested in animal experiments. Non-invasive BSIs would allow broadening this field of research, facilitating experimentation in healthy subjects and patients with motor disorders. Electroencephalography (EEG) constitutes the most common technique for non-invasive acquisition of brain signals. However, the low signal to noise ratio and artifacts often limit EEG applications. This problem aggravates when EEG is concurrently used with electromagnetic stimulation because it can contaminate the EEG signals and impede the estimation of cortical activity (Insausti-Delgado et al., 2020).

In the current study, we propose an innovative design for a non-invasive BSI, relying on the continuous EEG monitoring of brain activity, removal of stimulation artifacts by median filtering, and control of the trans-spinal magnetic stimulation (ts-MS) to volitionally (but artificially) contract lower limb muscles. The system was tested and validated in 10 healthy participants, with 4 different stimulation conditions. As a proof of the feasibility of this interface, we report the evaluation of different indicators: *(i)* the performance, robustness and decoding accuracy of the BSI, *(ii)* the usability and perception of all the users, and *(iii)* the neurophysiological effects of the system.

## 2 Materials and methods

### 2.1 Participants

Ten healthy participants (4 females, age = 29.5 ± 4.67 years) with no neurological disorders and complete mobility of lower limbs were recruited for the study. All the participants provided written informed consent before starting the experiment, which was approved by the Ethics Committee of the Faculty of Medicine of the University of Tübingen (Germany).

The participants were comfortably seated on a chair, with their back straight and their right leg slightly extended, having the knee and ankle joint angles around 120° and 90°, respectively. A wedge-shaped structure was used to place the foot and ensure the angle of the leg was kept constant (Figure 1a). Neurophysiological activity was recorded by electroencephalography (EEG) and electromyography (EMG).

**Figure 1.**
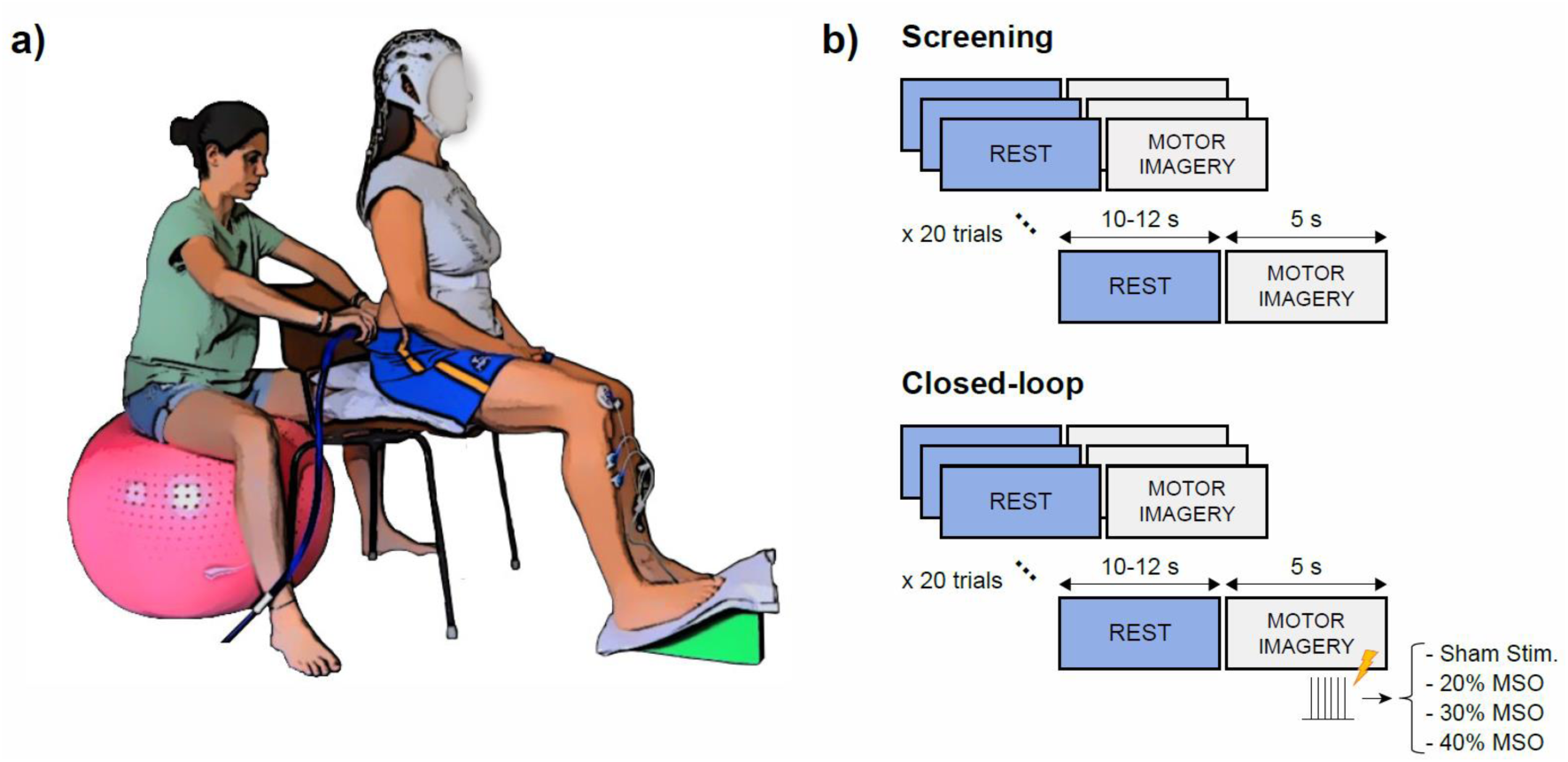
Experimental procedure. (a) Participant with the EEG system and EMG sensor on the right tibialis anterior (TA) muscle, and experimenter placing the coil for spinal stimulation. The experimenter in the picture is the first author of the paper and gives consent for publication of her image. (b) Block diagram of the two experiment phases: screening and closed-loop stimulation. Each phase included blocks of 20 trials, consisting of rest and motor imagery periods announced by an auditory cue of “Rest” and “Move”, respectively. During the closed-loop phase, contingent ts-MS was applied at 20 Hz when the motor imagery was detected from the EEG of the participant. Each closed-loop block was executed with a fixed intensity: 20% of the maximum stimulator output (MSO), 30% of the MSO, 40% of the MSO, or sham stimulation.

### 2.2 Experimental design and procedure

Each participant performed one session, including a screening phase and a closed-loop stimulation phase (Figure 1b). The screening consisted of 2 blocks of 20 trials each, which included rest (10-12 s) and motor imagery (5 s) periods, each announced by an auditory cue of “Rest” and “Move”, respectively. During rest, the participants were asked to relax and stay still without executing or imagining any movement. During motor imagery (MI), the participants were asked to perform kinesthetic motor imagery of the plantar dorsiflexion of the right leg (Neuper et al., 2005). The EEG data recorded during the screening was used to train a classifier to differentiate between the “rest” and “motor imagery” brain states.

The closed-loop stimulation phase consisted of 12 blocks of 20 trials each. We evaluated 4 stimulation conditions: 3 different ts-MS intensities and sham stimulation (further details in Section 2.4). We recorded 3 blocks of each condition, randomizing the sequence of intensities across subjects, resulting in 60 trials of each intensity. The timing of the stimulation trials was identical to the screening trials. Closed-loop feedback was given according to the decoded brain patterns during the MI periods (note that the stimulation was off during resting periods).

### 2.3 Data acquisition

EEG activity was recorded with a commercial Acticap system (BrainProducts GmbH, Germany), with 32 channels placed on FP1, FP2, F7, F3, F4, F8, FC3, FC1, FCz, FC2, FC4, C5, C3, C1, Cz, C2, C4, C6, CP3, CP1, CPz, CP2, CP4, T7, T8, P7, P3, Pz, P4, P8, O1, and O2, following the international 10/20 system (Seeck et al., 2017). The ground and reference electrodes were located at FPz and Fz, respectively. The recording electrodes were connected to a monopolar BrainAmp amplifier (BrainProducts GmbH, Germany).

EMG activity from the right tibialis anterior (TA) muscle was recorded using Ag/AgCl bipolar electrodes (Myotronics-Noromed, Tukwila, Wa, USA) combined with an MR-compatible BrainAmp amplifier (BrainProducts GmbH, Germany). The recording electrodes had an inter-electrode space of 4 cm. The ground electrode was placed on the right patella. All the signals were synchronously acquired at 1 kHz sampling rate.

### 2.4 Trans-spinal magnetic stimulation (ts-MS)

Due to the unnatural (and occasionally uncomfortable) sensation that the participants can experience with ts-MS, and to ensure that they were able to bear the stimulation, a familiarization session was conducted with all of them on a separate day before the experiment. We used a magnetic stimulator (Magstim Rapid2, Magstim Ltd, UK) with a circular coil (Magstim 90 mm Coil, Magstim Ltd, UK) to provide the ts-MS (biphasic single cosine cycle pulses of 400 µs).

Before starting the recording, we localized and marked the vertebrae from T12 to L5 according to anatomical landmarks. The circular coil was initially centered over the midline of the intervertebral space of T12 and shifted towards L5, advancing one vertebra in each step (coil currents directed clockwise). Single pulse stimulation was delivered above the motor threshold to locate the hot-spot of the TA muscle (i.e., the spot that led to the largest trans-spinal motor evoked potentials in 10 trials). This spot was marked for the closed-loop stimulation phase.

During the closed-loop phase, continuous brain-controlled ts-MS was applied at 20 Hz (Sasada et al., 2014). According to our previous experimental evidence, the spinal motor threshold (i.e., the minimum intensity needed for eliciting at least 5 motor evoked potentials out of 10 trials with at least 50 µV of peak-to-peak amplitude) lays between 25% and 40% of the maximum stimulator output (MSO) (Insausti-Delgado et al., 2019, 2018). According to these values, we defined 4 conditions for the closed-loop stimulation: *(i)* ts-MS at 20% of the MSO, *(ii)* ts-MS at 30% of the MSO, *(iii)* ts-MS at 40% of the MSO, and *(iv)* sham stimulation. For the sham stimulation, the experimenter held the coil 1 m away from the participant, so that the stimulation took place and the participants had auditory but no sensory feedback. In this sham condition, the stimulation was set to 30% of the MSO. For the three real ts-MS conditions, the coil was placed on the hot-spot.

### 2.5 Detection of movement intention

After the 2 screening blocks, a classifier was trained to discriminate between the brain states of rest and MI.

#### 2.5.1 Data preprocessing

The EEG data were band-pass filtered with a 4^th^ order Butterworth filter between 1 and 50 Hz. The signals were trimmed down to 15-second trials (from -10 s to +5 s with respect to the MI cue), and subsampled to 100 Hz. Optimized spatial filtering (OSF) was applied to improve the estimation of the task-related motor cortex activation. We considered 17 electrodes (from the 32 recorded) to measure this activation: FC3, FC1, FCz, FC2, FC4, C5, C3, C1, Cz, C2, C4, C6, CP3, CP1, CPz, CP2 and, CP4. The signal of these electrodes was band-pass filtered between 7 and 15 Hz (4^th^ order Butterworth), to isolate the modulation of the alpha rhythm (López-Larraz et al., 2014). The OSF calibration consists of a gradient-descent optimization to find weights for the linear combination of electrodes that minimizes alpha power during MI and maximizes it during rest. This has been validated as an effective automated method to improve the measurement of event-related desynchronization (ERD) of sensorimotor activity during motor tasks (Graimann and Pfurtscheller, 2006). The result of this process is a virtual channel that synthesizes the activation over the motor cortex. The OSF weights were computed using the trials of the screening phase and kept fixed during the closed-loop phase.

#### 2.5.2 Feature extraction

A one-second sliding window, with 200 ms sliding-step, was applied to each 15-second trial of the OSF virtual channel in the interval [-3, -1] s for the rest class and [1, 3] s for the MI class (i.e., 6 windows per class and trial). The power spectrum between 1 Hz and 50 Hz was calculated for each of these windows using a 20^th^-order autoregressive (AR) model with 1 Hz resolution, based on the Burg algorithm (Burg, 1967). The most discriminant range of frequencies to separate between rest and MI classes was selected by visually inspecting the signed r-squared values (point-biserial correlation coefficients). Despite alpha ([7-15] Hz) and beta ([15-30] Hz) being generally the most reactive frequency bands during MI, we restricted the selection of features to the alpha range only, to avoid the repetitive ts-MS at 20 Hz interfering with our brain features of interest. The power values within the selected frequency range were averaged, resulting in one unique feature.

All the extracted windows from the screening trials, transformed into one feature per window, were z-score normalized and fed to a linear discriminant analysis (LDA) classifier to distinguish between both classes.

#### 2.5.3 Classification

During the closed-loop blocks, the classifier analyzed the EEG activity in real-time and activated the stimulator when the MI brain states were detected. A new block of EEG data arrived every 200 ms. This block was median-filtered (see details in 2.5.3.1), band-pass filtered between 1 and 50 Hz, OSF filtered (using the coefficients computed from the screening data), and appended to a one-second ring-buffer to compute the newest power output (following 2.5.2). The classifier determined whether this power output corresponded to rest or MI class and triggered ts-MS as long as MI was detected, providing continuous feedback. Note that the stimulation was deactivated during the rest periods, avoiding stimulation due to false positives.

To deal with EEG-nonstationarities and potential changes of cortical activation patterns due to ts-MS, we continuously updated the normalization coefficients of the features (initially, the mean and standard deviation of the training dataset). We kept two 48-second buffers, one for rest and one for MI, with the most recent features of these classes. The mean and standard deviation of these two buffers, concatenated together, were used as the normalization coefficients in each iteration before passing the feature vector to the classifier.

##### 2.5.3.1. Median filtering for ts-MS contamination removal

Using electromagnetic currents for stimulating the nervous system can introduce undesirable noise to the neural activity. As characterized in (Insausti-Delgado et al., 2017), the ts-MS distorts the EEG recordings, introducing peaks of short duration (∼10 ms) and large magnitude. A median filter is a suitable method for minimizing the influence of the ts-MS contamination in the EEG signal (Insausti-Delgado et al., 2017, 2020). We applied the median filter as a sliding window of 20 ms in one-sample steps, calculating the median value for each window. This filter attenuates these large peaks preserving the sensorimotor oscillatory activity. A detailed characterization of how this filter can be used to remove similar peaks due to electrical stimulation can be found in (Insausti-Delgado et al., 2020).

### 2.6 Decoding accuracy

The performance of the classifier for each participant was estimated in terms of average decoding accuracy, calculated as the mean of the true positive rate (TPR) and the true negative rate (TNR). The TPR quantifies the success of the classifier during the MI period, defined as the time interval [1, 4] s. The TNR measures the classifier success during the rest period, which was defined as the time interval [-4, -1] s.

### 2.7 Neurophysiological measurements

Our BSI has been devised to be used as a rehabilitative tool for patients that have motor impairments. For future interventions based on BSIs, the potential of these systems to interact with cortico-spinal and spino-muscular circuitry is a relevant aspect. With the data recorded during the closed-loop blocks, we conducted some neurophysiological measures to assess the interactions of the BSI with the nervous system.

#### 2.7.1 Trans-spinal motor evoked potentials (ts-MEP)

Ts-MS can activate the peripheral nervous system, exciting the spinal nerves at their exit through the intervertebral foramina towards the muscles, resulting in ts-MEPs (Matsumoto et al., 2013; Ugawa et al., 1989). The recruitment of peripheral motor nerves was demonstrated by measuring the ts-MEPs at the tibialis anterior (TA) muscle during spinal stimulation. EMG signals were high-pass filtered at 10 Hz with a 4^th^ order Butterworth filter and trimmed down to 45-ms epochs (from -5 to 40 ms with respect to the stimulation pulse). Epochs corresponding to the same ts-MS intensity were pooled together and averaged for each subject. The peak-to-peak amplitude of the ts-MEPs for each intensity was determined as the difference between the maximum and the minimum values of the averaged potential.

#### 2.7.2 Trans-spinal somatosensory evoked potentials (ts-SEP)

Spinal stimulation can activate the sensory cortex via the ascending pathways from the spine, which can be quantified as ts-SEPs (Kunesch et al., 1993). The EEG activity of the CPz channel during closed-loop was high-pass filtered using a 4^th^ order Butterworth filter at 3 Hz. Signals were aligned to the stimulation artifact and epoched to 45-ms periods (from - 5 to 40 ms with respect to the ts-MS pulse). Epochs were grouped according to their intensity and averaged for each subject. The peak-to-peak amplitude of the averaged ts-SEP was calculated as the difference between the maximum and the minimum values for each ts-MS intensity.

### 2.8 Usability assessments

The participants were asked to evaluate the degree of pain, discomfort and concentration at the end of each closed-loop block. They had to grade between 0 (very low) and 10 (very high): *(i)* how painful the stimulation was, *(ii)* how uncomfortable the stimulation was, and *(iii)* how easy it was to perform the motor imagery while being stimulated.

### 2.9 Statistical analysis

We studied the effect of stimulation condition on the BSI performance and on the neurophysiological measurements. The Shapiro-Wilk test was used to determine the Gaussianity of the data. To assess the effect of stimulation on MI decoding accuracy, we used a repeated measures analysis of variance (ANOVA), with stimulation condition as factor (4 levels: ts-MS at 20% of the MSO, ts-MS at 30% of the MSO, ts-MS at 40% of the MSO, and sham stimulation) and decoding accuracy as dependent variable. Post-hoc comparisons were conducted using paired t-tests with Bonferroni correction. To evaluate the influence of the median filter on the BSI performance, we ran paired t-tests with filtering as factor (with and without median filter) and decoding accuracy as dependent variable for each stimulation condition. To study the influence of stimulation intensity on the peak-to-peak amplitude of ts-MEPs and ts-SEPs we used Friedman’s test (3 levels: ts-MS at 20% of the MSO, ts-MS at 30% of the MSO, ts-MS at 40% of the MSO). Paired post-hoc comparisons were performed using the Wilcoxon signed-rank test to analyze significant amplitude differences between intensity pairs. All the statistical tests were conducted in IBM SPSS 25.0 Statistics software (SPSS Inc., Chicago, IL, USA).

## 3 Results

### 3.1 Brain-spine interface control

Seven out of the ten subjects showed strong cortical activation patterns during MI in the screening data, revealed as a significant event-related desynchronization (ERD) in alpha and beta bands (Figure 2).

**Figure 2.**
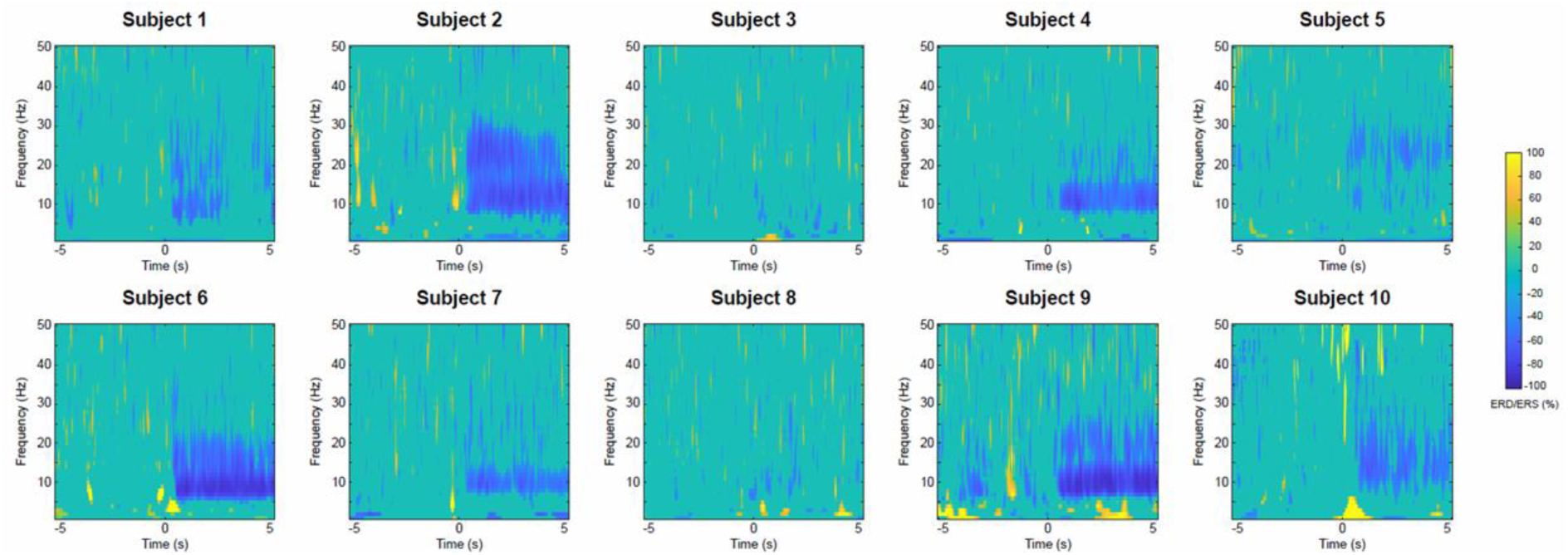
Cortical activation during motor imagery (computed from the screening blocks). Time-frequency maps for each individual, representing the event-related (de)synchronization ERD/ERS (Pfurtscheller and Lopes da Silva, 1999) of the optimized spatial filter (OSF) channel. Time 0 s represents the auditory cue to start the motor imagery.

Video 1 shows one representative participant controlling the BSI. The participant was asked to rest or to perform MI of the right ankle dorsiflexion, guided by auditory cues. The EEG activity was processed (median filtered, band-pass filtered and OSF filtered) in real-time. To prove the efficacy of the system to remove online stimulation artifacts, the activity of the OSF channel with and without median filtering is displayed. The classifier triggered the ts-MS when the MI brain states were detected. Note that the stimulation was off during resting periods. The experimenters in the video are the authors of the paper and give consent for publication of their video.

The average decoding accuracies for all the participants were 61.7%, 57.9%, 58.4% and 59.6% for sham stimulation, ts-MS at 20% of the MSO, ts-MS at 30% of the MSO and ts-MS at 40% of the MSO, respectively (Figure 3 top panel). There was no significant effect of stimulation condition on decoding accuracy, as revealed by the repeated measures ANOVA (F(3, 27) = 1.912, *p* = 0.151). As a post-hoc analysis, we discarded the data of the three participants who did not show a modulation of the sensorimotor alpha rhythm during MI in the screening phase (S3, S5 and S8). These three participants had a decoding accuracy around chance level in the closed-loop phase for every stimulation condition. When we excluded them from the analysis, the average decoding accuracy increased up to 66.2%, 61.8%, 63.5% and 66.1%, respectively (Figure 3 bottom panel). These values were not significantly different between ts-MS intensities either (repeated measures ANOVA, F(3, 18) = 2.008, *p* = 0.149).

**Figure 3.**
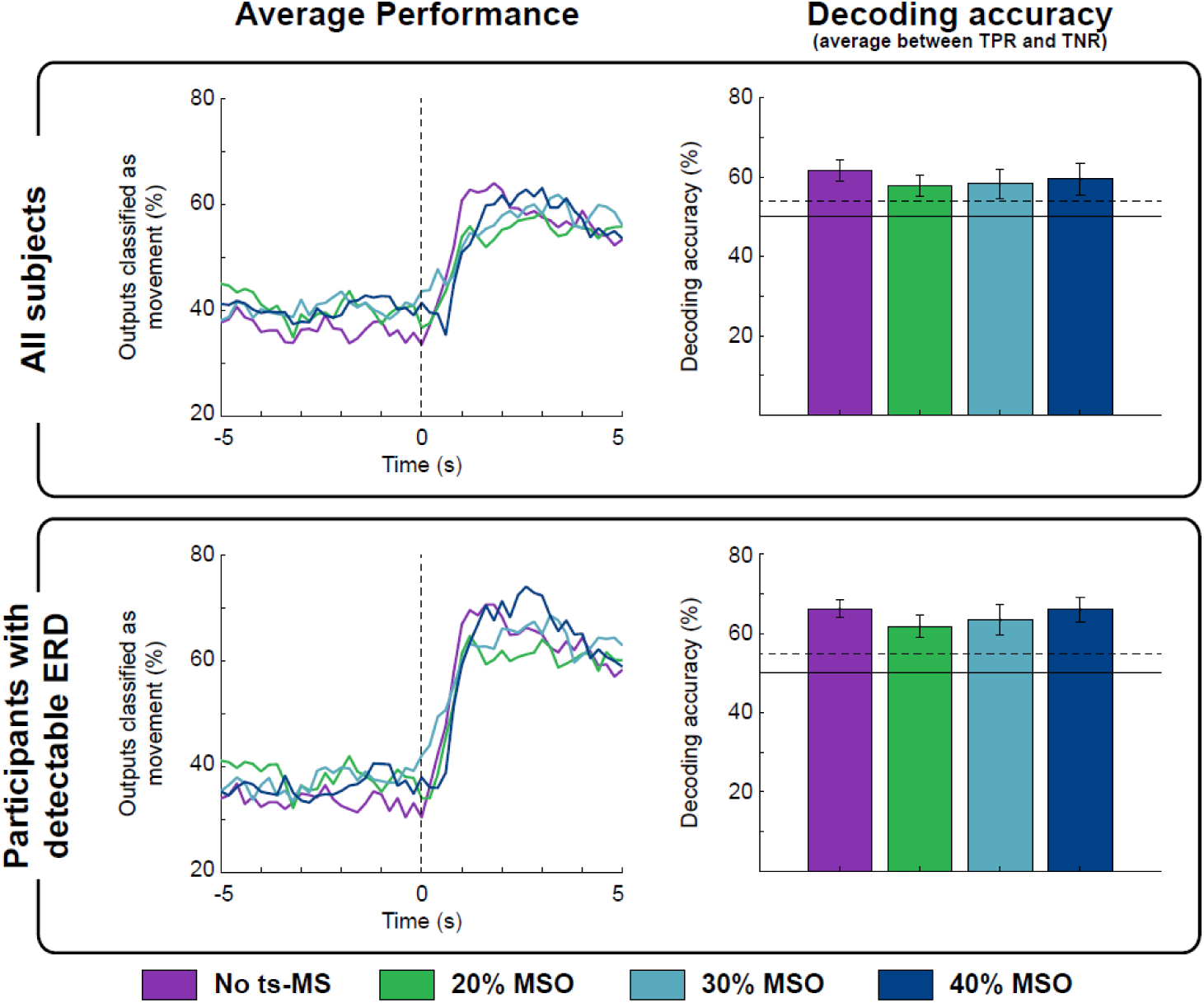
Average response of the classifier for all the stimulation conditions. (Left) Average time-response of the classifier for each stimulation condition. Each line represents the percentage of the outputs classified as motor imagery, averaged over all the participants. Notice that time 0 s is the beginning of the motor imagery period, and outputs prior to t=0 represent false positives, while outputs after t=0 mean true positives. (Right) Decoding accuracy, calculated as the mean between true negative rate (TNR) in the time interval [-1, -4] s and true positive rate (TPR) in the time interval [1, 4] s. The dashed line shows the confidence interval of the chance level (alpha = 0.05), calculated on the basis of all the test trials, according to (Müller-Putz et al., 2008). Panels show the values averaged for all the participants (top) and for the seven participants with detectable MI-related desynchronization in the alpha frequency band (bottom).

### 3.2 Effect of artifact removal

We also studied how ts-MS affects the cortical activity and the performance of the BSI. Stimulation artifacts distort the ongoing EEG activity, hindering its processing. The median filter effectively eliminated the high-amplitude peaks, allowing the quantification of sensorimotor modulation (Figure 4a). Figure 4b displays the estimated cortical activity of a representative participant during closed-loop ts-MS at 40% of the MSO (i.e., the highest stimulation intensity) with and without applying the median filtering. If the stimulation artifacts are not eliminated, they are observable in the EEG as a broadband event-related synchronization (ERS), or power increase, covering the frequencies of interest (Figure 4b left). Median filtering revealed the significant event-related desynchronization (ERD) of alpha and beta frequencies (Figure 4b right), which allowed the classifier to decode the MI (Figure 4c). We calculated the decoding accuracy of MI with and without median filter for each stimulation condition for those participants with detectable ERD. Paired t-tests revealed that applying the median filter leaded to significantly higher decoding accuracies in closed-loop ts-MS at 20% (*t*(6) = -3.948, *p* = 0.008), ts-MS at 30% (*t*(6) = -5.079, *p* = 0.002) and ts-MS at 40% of the MSO (*t*(6) = -6.331, *p* = 0.001) (Figure 4d).

**Figure 4.**
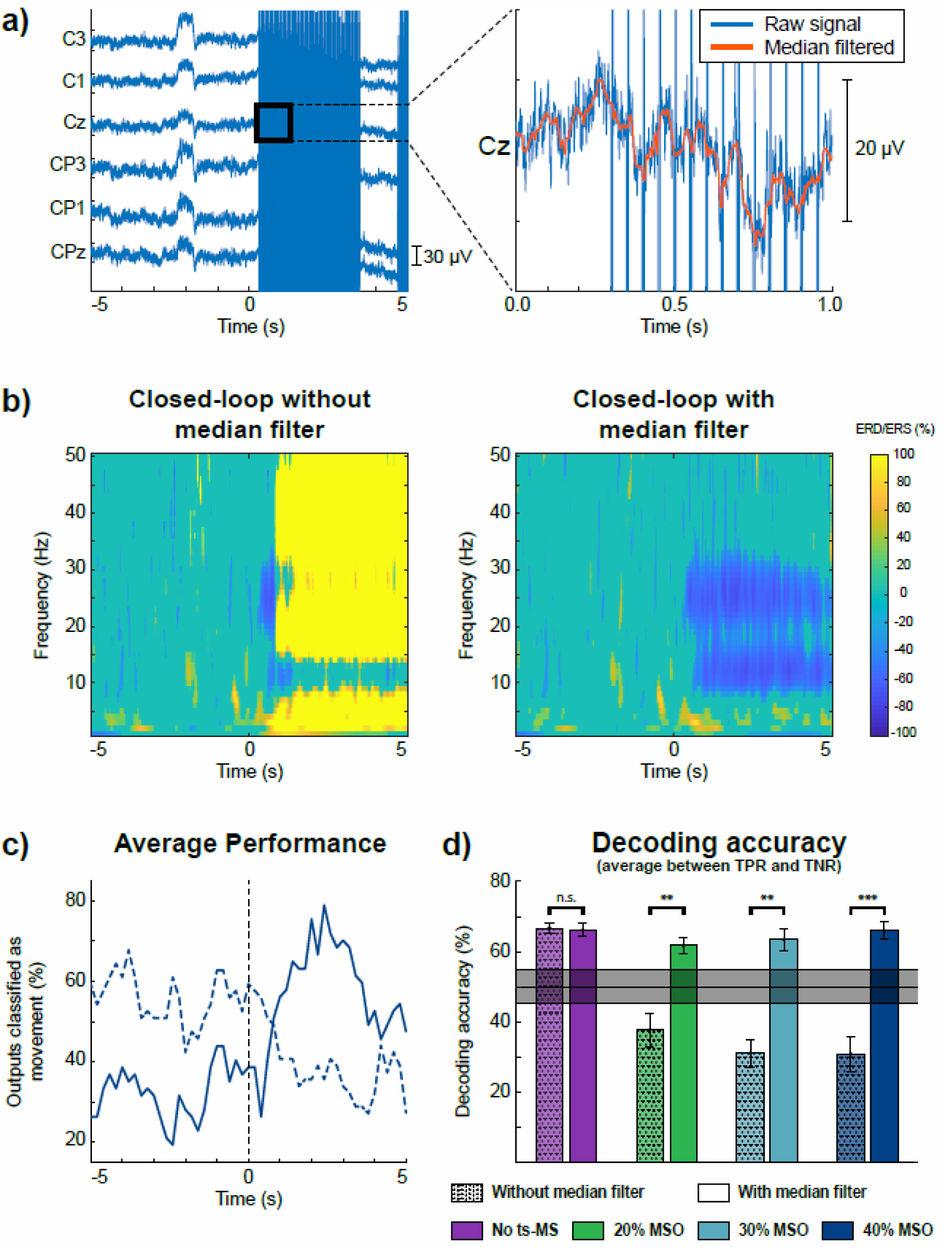
Characterization of stimulation artifacts and their effects on cortical activity and decoding accuracy. The results for one representative participant are displayed in the most unfavorable scenario, with stimulation at 40% of the MSO. (a) EEG trace of one trial, showing the effect of the stimulation as high-amplitude artifacts. Zooming into a one-second segment, the details of the signal without (blue) or with (orange) median filtering can be appreciated. (b) Grand average time-frequency maps without (left) and with (right) median filter. Time 0 s corresponds to the onset of the motor imagery. (c) Average time-response of the classifier without (dashed) and with (solid) median filter. (d) Decoding accuracy calculated as the mean between TNR and TPR for each stimulation condition with and without median filter for the seven participants with detectable MI-related desynchronization in the alpha frequency band. The shaded gray area shows the confidence interval of the chance level (alpha = 0.05), calculated on the basis of all the test trials, according to (Müller-Putz et al., 2008). The asterisks indicate significant differences (** for p >0.01, *** for p >0.001) between decoding accuracies when median filtered is applied.

### 3.3 Neurophysiological analysis

#### 3.3.1 Trans-spinal motor evoked potentials (ts-MEP)

By analyzing the ts-MEPs, we assessed the neurophysiological effects of the stimulation on the peripheral nervous system (Figure 5a). We extracted the ts-MEPs from the closed-loop stimulation blocks (Figure 5b), and compared their amplitude according to the stimulation intensity (Figure 5c). The peak-to-peak amplitude of the ts-MEPs was significantly affected by the intensity used (Friedman’s χ^2^(2) = 15.80; p < 0.001). Post-hoc comparisons revealed significantly larger ts-MEPs amplitudes at 40% of the MSO compared to 20% (Z = -2.803, p = 0.015) and compared to 30% (Z = -2.803, p = 0.015) (Figure 5d).

**Figure 5.**
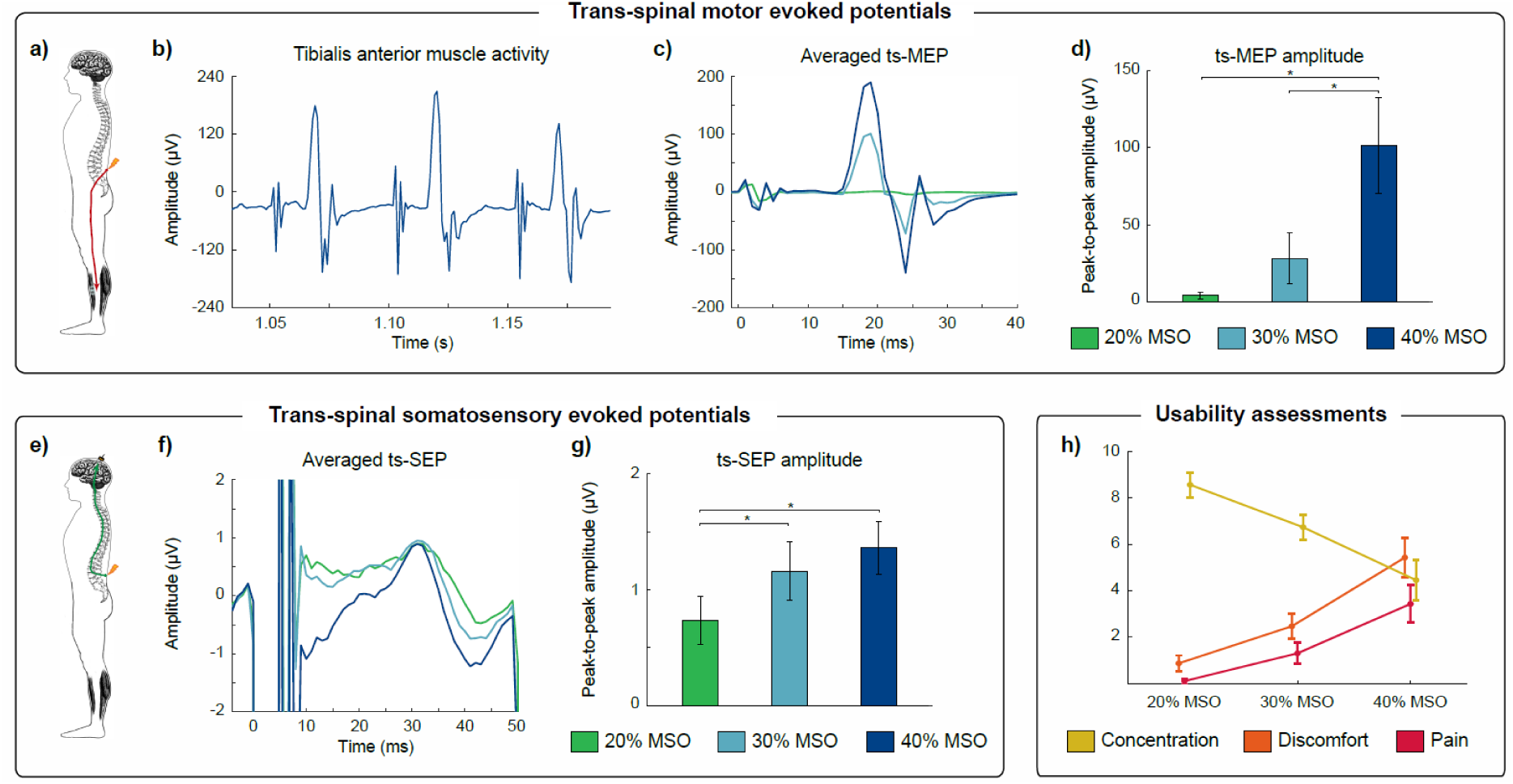
Neurophysiological and usability assessments. Upper panel, a) Recruitment of efferent pathways from the spine to the tibialis anterior (TA) muscle by ts-MS. b) EMG trace of the TA muscle of a representative participant during closed-loop stimulation at 40% ts-MS. The artifacts appear approximately at t = 1.05, 1.10 and 1.15 s (20 Hz stimulation), while the ts-MEPs are induced ∼15 ms after the stimulation (peripheral motor conduction time from the spine to the TA). c) Average ts-MEPs of a representative participant for the three different stimulation intensities. d) Mean peak-to-peak amplitude of ts-MEPs and standard errors averaged over all participants for each stimulation intensity. The asterisks indicate significant (p < 0.05, Bonferroni corrected) differences in ts-MEP amplitude between ts-MS intensities. Bottom-left panel, e) Recruitment of afferent pathways from the spine to the cortex via ts-MS. f) Average ts-SEPs of a representative participant for the three different stimulation intensities. g) Mean peak-to-peak amplitude of ts-SEPs and standard errors averaged over all participants for each stimulation intensity. The asterisks indicate significant (p < 0.05, Bonferroni corrected) differences in ts-SEP amplitude between ts-MS intensities. Bottom-right panel, h) Usability scores for concentration, discomfort and pain, averaged over all participants.

#### 3.3.2 Trans-spinal somatosensory evoked potentials (ts-SEP)

We computed the ts-SEPs to assess the neurophysiological effects of the stimulation on the central nervous system (Figure 5e). As for the ts-MEPs, we averaged the ts-SEPs for each subject, grouping by stimulation intensity. The ts-MS produced a positive peak with a latency of 30 ms and a negative peak at 40 ms (Figure 5f). The peak-to-peak amplitude of the ts-SEP was significantly affected by the stimulation intensity (Friedman’s χ^2^(2) = 11.40; p = 0.003). Post-hoc paired comparisons showed significantly smaller ts-SEPs when stimulating at 20% compared to 30% (Z = -2.599, p = 0.027) or 40% (Z = -2.701, p = 0.021) of the MSO (Figure 5g).

### 3.4 Usability assessments

Our descriptive analysis on usability shows that the participants perceived more discomfort and pain, and decreased concentration on the MI task, as the intensity of ts-MS increased (Figure 5h). All the participants described the stimulation at higher intensities as uncomfortable, rather than painful. No adverse effects due to ts-MS were reported by, or observed in, any of the participants.

## 4 Discussion

In this paper, we report on the first non-invasive brain-spine interface (BSI), based on the continuous control of trans-spinal magnetic stimulation (ts-MS) guided by EEG. Our BSI enables the direct association of cortical activity encoding motor intentions with the activation of afferent (from the spine to the somatosensory cortex) and efferent (from the spine to the lower-limb muscles) pathways. This natural approach to link brain activity with the peripheral nervous system could be used to exploit neuromodulatory mechanisms and might constitute a relevant tool for rehabilitation of patients with paralysis. The here presented findings provide sufficient basis towards designing, developing and further evaluating this innovative approach.

Brain-controlled spinal cord stimulation can potentially be employed for assistive or rehabilitative purposes by patients with lower-limb paralysis. Spinal cord stimulation has been used to neuromodulate the spinal circuitry, supporting motor recovery after lower-limb paralysis (Edgerton and Roy, 2012; Nardone et al., 2015b; Taccola et al., 2018). The first studies in animals evidenced the neuromodulatory properties of dorsal root stimulation of the spine (Budakova, 1972; Grillner and Zangger, 1979). Later investigations using epidural spinal stimulation supported previous findings and demonstrated the capacity of stimulation to enable standing and gait in paralyzed rodents and cats (Gerasimenko et al., 2008; Ichiyama et al., 2005; Wenger et al., 2016). In humans, epidural stimulation of the spinal cord has been shown to facilitate locomotor-like patterns and produce long-lasting motor recovery after intensive training in spinal cord injury patients (Angeli et al., 2014; Gill et al., 2018; Grahn et al., 2017). However, controlling and modulating the stimulation based on brain activation is a more natural approach than continuously stimulating the spinal circuits (McPherson et al., 2015). In fact, the contingent association of cortical activity produced by the intention to move a paralyzed limb and the afferent volley generated by spinal stimulation can exploit Hebbian mechanisms and facilitate functional recovery (Bonizzato et al., 2018; Mrachacz-Kersting et al., 2016; Ramos-Murguialday et al., 2013). This association is the basic operating principle of brain-spine interfaces (Alam et al., 2016).

Brain-spine interfaces developed to date involve implantable technologies and have only been tested in animal models (Alam et al., 2014; Bonizzato et al., 2018; Capogrosso et al., 2016; McPherson et al., 2015; Nishimura et al., 2013b). Compared with continuous spinal stimulation, brain-controlled stimulation has been shown to enhance stepping quality and accelerate locomotor recovery (Bonizzato et al., 2018; Capogrosso et al., 2016). In humans, the only approaches presenting closed-loop volitional control of spinal stimulation proposed non-brain-commanded paradigms. On one hand, Nishimura and colleagues proposed the use of EMG of the arm to control non-invasive magnetic spinal stimulation in healthy subjects and SCI patients (Nakao et al., 2015; Sasada et al., 2014). On the other hand, Courtine and colleagues implanted epidural electrical stimulation electrodes in the lumbar spinal cord of SCI patients and used inertial measurement units (IMUs) located on the feet to control the stimulation (Wagner et al., 2018). Our approach relied on extracting motor commands non-invasively from brain activity by EEG to provide closed-loop control of transcutaneous magnetic stimulation of the spinal cord.

Non-invasive brain-machine interfaces (BMIs) allow the transmission of volitional cortical commands to control rehabilitative devices (López-Larraz et al., 2018b; Millán et al., 2010; Wolpaw et al., 2002). For instance, there is ample evidence demonstrating contingent EEG control of robotic exoskeletons with patients (Ang et al., 2015; Ramos-Murguialday et al., 2013). Electric and magnetic neurostimulation can also be integrated with non-invasive BMIs. However, to date, contingent online control of such neurostimulators has not been achieved, since the stimulation introduces currents to the body, causing strong artifacts that hinder extracting reliable information from the recordings of brain activity. Therefore, BMIs integrating neurostimulation have only been proposed triggering predefined stimulation patterns, not allowing a continuous control nor contingency (Biasiucci et al., 2018; Osuagwu et al., 2016; Pfurtscheller et al., 2005; Trincado-Alonso et al., 2017).

Dealing with stimulation artifacts is a challenge for closed-loop neural interfaces. Different approaches have been proposed for cleaning stimulation contamination from invasive and non-invasive neural recordings, such as blanking, interpolation or linear regression reference (LRR) (Iturrate et al., 2018; Walter et al., 2012; Young et al., 2018). For invasive recordings, regression methods have been proven effective to eliminate the stimulation artifact (Young et al., 2018), mainly due to the low inter-electrode impedance variability and within-session stability, allowing closed-loop neurostimulation (Ajiboye et al., 2017; Bouton et al., 2016). However, none of these methods has been proven effective for EEG recordings, and the estimation of cortical activation during stimulation is biased even if blanking or interpolation of the artifacts is used (Walter et al., 2012).

We proposed the use of a median filter, since it can eliminate high-amplitude peaks in a time-series without causing signal discontinuities (which is the main problem of blanking or interpolation). Its main limitation is that it attenuates the activity at higher frequencies (exponential attenuation between 0 Hz and the 1/ws Hz, with ws being the length of the window of the median filter) (Insausti-Delgado et al., 2020). However, this frequency-dependent attenuation does not have a big impact on the sensorimotor alpha oscillations (7-15 Hz) that we used to detect the motor imagery. To the best of our knowledge, this is the first time a non-invasive closed-loop system controls the stimulation in real-time and effectively deals with these artifacts.

Our adaptive decoder successfully dealt with the changes in brain activity due to ts-MS. The accuracy of the decoder was within the level of acceptance for closed-loop rehabilitative neuroprosthetics (Ramos-Murguialday et al., 2013). Remarkably, we demonstrated that the accuracy was not influenced by the stimulation intensity, which allows the implementation of different stimulation protocols, such as above or below motor threshold (considering that the motor threshold of healthy subjects is between 25% and 40% of the MSO). All participants reported a decrease in usability of the system with higher stimulation intensities (i.e., more pain and discomfort, and less ability to concentrate on the task). However, their subjective perception of usability at high intensity did not affect the performance of the BSI.

The next natural step following this study would be to demonstrate the efficacy of the brain-controlled stimulation to promote Hebbian mechanisms that induce neuroplastic changes in human nervous system, as it has already been proven invasively in rodents and primates (Bonizzato et al., 2018; Capogrosso et al., 2018, 2016; McPherson et al., 2015). An exhaustive battery of assessments should be conducted, including the evaluation of motor and sensory pathways and spinal neural processes, to characterize in detail the neurophysiological effects of a BSI-based intervention. Although we did not conduct assessments of synaptic efficacy in this preliminary analysis, we studied the neuromodulatory effects during closed-loop stimulation. We demonstrated the capability of our platform to engage the central and peripheral nervous system as expected. The ts-MS activated efferent pathways, inducing ts-MEPs in lower limb muscles, and afferent pathways, producing ts-SEPs. These findings prove that both afferent and efferent neuromodulation are intensity dependent, confirming previous results (Matsumoto et al., 2013; Rossini et al., 2015).

In conclusion, in this study we have proposed and validated the first non-invasive brain-spine interface. Further research should focus on studying the feasibility of this system as a rehabilitative tool in patient populations such as SCI or motor stroke. The role of ts-MS parameters (i.e., frequency, intensity, dose, etc.) on the excitability of spinal neural networks should also be disclosed in future investigations. Computational modelling might be a key tool to understand the ongoing mechanisms involved in spinal neuromodulation due to ts-MS in order to optimize interventions based on spinal cord stimulation that enhance functional recovery (Formento et al., 2018; Greiner et al., 2020; Khadka et al., 2020). Nevertheless, the here presented findings constitute the first steps towards the application of non-invasive BSIs as a novel neuroscientific and therapeutic tool.

## 5 Acknowledgments

This study was funded by the Bundesministerium für Bildung und Forschung BMBF MOTORBIC (FKZ 13GW0053) and AMORSA (FKZ 16SV7754), and the Fortüne-Program of the University of Tübingen (2422-0-1 and 2556-0-0 to ELL, and 2452-0-0 to ARM). The work of AID was funded by the Basque Government’s scholarship for predoctoral students.

